# Metabolic exploitation of the bacterial arginine deiminase pathway extends *Caenorhabditis elegans* healthspan

**DOI:** 10.64898/2026.09.07.749978

**Authors:** Arisa Ito, Daisuke Kyoui, Taketo Kawarai, Hiroshi Matsufuji, Chise Suzuki

## Abstract

While certain lactic acid bacteria extend *Caenorhabditis elegans* lifespan, the underlying mechanisms remain obscure. We investigated the lifespan-extending effects of *Lactococcus lactis* JCM 5805 (Lc5805). While live and heat-treated Lc5805 both extended mean lifespan, live Lc5805 uniquely prolonged maximum lifespan to 25 days, compared with 21 days for control and heat-treated groups. Ingesting live Lc5805 upregulated 74 cuticle-formation genes, predominantly collagen genes, supporting structural integrity without activating host stress responses. Metabolomic analyses revealed that live Lc5805 consumption induced L-citrulline and L-ornithine accumulation in *C. elegans*, which lacks a functional urea cycle. Intestinal expression of bacterial arginine deiminase (*arcA*), contrasted with the absence of citrulline accumulation in ADI-lacking bacteria and heat-treated groups, indicates that citrulline accumulation requires active bacterial enzymes to operate the arginine deiminase pathway in the gut. By utilizing bacterial metabolites and enzymes to support collagen maintenance, the host avoids endogenous energy expenditure via “metabolic exploitation.”

## Introduction

*Caenorhabditis elegans* is an excellent model organism for studying aging and metabolism due to its short lifespan and well-characterized metabolic pathways (Consortium 1998; Giunti *et al*. 2021). In laboratory settings, *C. elegans* is conventionally fed *Escherichia coli* OP50; however, in its natural habitat, it primarily feeds on live bacteria in decaying plant matter (Félix and Duveau 2012).

Unlike vertebrates, which detoxify toxic ammonia into less harmful nitrogenous molecules such as urea or uric acid prior to excretion, *C. elegans* directly excretes ammonia. Consequently, *C. elegans* lacks four key genes required for arginine (Arg) biosynthesis via the urea cycle—carbamoyl phosphate synthetase 1 (*CPS1*), ornithine transcarbamylase (*OTC*), argininosuccinate synthase 1 (*ASS1*), and argininosuccinate lyase (*ASL*)—making Arg an essential amino acid for this organism (Zecic *et al*. 2019). In mammals, the urea cycle serves not only to convert toxic ammonia into harmless urea, but its metabolic intermediates are also intrinsically linked to the synthesis of major amino acids constituting collagen. Specifically, ornithine is converted into glutamate-5-semialdehyde, which subsequently serves as a precursor for proline, a key component of collagen (Albaugh *et al*. 2017; Barbul 2008). Thus, a stable supply of proline from ornithine via the urea cycle, which serves as a precursor for hydroxyproline, is essential for mammals to synthesize and maintain collagen properly. However, *C. elegans* has evolutionarily relinquished the energy-costly urea cycle required for ammonia disposal. Instead, it relies on an alternative pathway from glutamate to supply the proline necessary for collagen production.

Furthermore, *C. elegans* lacks nitric oxide synthase (NOS), which synthesizes nitric oxide (NO) from Arg. Interestingly, feeding *C. elegans* with *Bacillus* strains possessing functional NOS extends its lifespan and enhances its stress tolerance (Gusarov *et al*. 2013). This suggests that *C. elegans* compensates for its own lack of NO production by “hijacking” NO generated by its naturally occurring dietary bacteria.

In addition to bacterial NO production, recent studies have highlighted the importance of host collagen and amino acid metabolism in lifespan extension and host–microbiome interactions. Collagen gene expression has been directly implicated in the regulation of aging and longevity in *C. elegans* (Wang *et al*. 2026). Specifically, the neuronal G-protein-coupled receptor NPR-8 orchestrates lifespan shortening at high temperatures and lifespan extension at low temperatures by neuroendocrinely regulating collagen genes via the transcription factor DAF-16. Because increased collagen expression is a hallmark common to various longevity interventions and stress resistance enhancements, maintaining collagen expression appears to be vital for healthy aging (Palani *et al*. 2023). Moreover, supplementation with individual amino acids—most notably serine and proline—has been shown to extend lifespan, with the exceptions of aspartate and phenylalanine (Edwards *et al*. 2015). Similarly, single-agent supplementation with L-citrulline (Cit) confers beneficial effects on healthspan in *C. elegans* (Rajcic *et al*. 2023).

Ingestion of specific lactic acid bacteria (LAB) has also been reported to markedly extend the lifespan of *C. elegans* (Kishimoto *et al*. 2024; Komura *et al*. 2024; Saito *et al*. 2026). However, the exact molecular mechanisms underlying these effects, as well as the host metabolic responses (e.g., changes in amino acid metabolism or collagen homeostasis) induced by LAB consumption, remain largely a “black box.”

In this study, we performed comprehensive gene expression and metabolite profiling (transcriptomic and metabolomic analyses) in *C. elegans* fed probiotic strains, namely *Lactococcus lactis* JCM 5805^T^ and *Lactiplantibacillus plantarum* MAFF 401534. Through this approach, we aimed to elucidate the real-time metabolic interactions between the host and ingested LAB.

## Materials and methods

### Bacterial strains and culture condition

Bacterial strains used in the present study were as follows: *E. coli* OP50 (OP50) was provided from Prof. Shohei Mitani of Tokyo Women’s Medical University. *L. plantarum* MAFF 401534 (Lp1534) and *L. lactis* subsp. *lactis* MAFF 400114 (Lc114) were purchased from The Research Center of Genetic Resources, NARO (National Agriculture and Food Research Organization). *L. lactis* G50 (LcG50) was provided from NARO. *L. lactis* JCM 5805 (Lc5805) was purchased from the JCM (RIKEN BRC, Tsukuba, Japan). OP50 was cultured overnight in LB medium at 37°C. Lp1534 and Lc5805 were cultured overnight in MRS medium at 30°C, while Lc114 and LcG50 were cultured overnight at 30°C in M17 medium supplemented with 0.5% glucose (hereafter referred to as GM17). After harvesting, the bacterial cells were washed with M9 buffer and adjusted to a concentration of 100 mg/mL. Heat treatment was performed by incubating 100 μL of the bacterial suspension in a heat block at 95°C for 10 min.

### *Caenorhabditis elegans* strains and growth conditions

*C. elegans* Bristol strain N2 was provided from Prof. Shohei Mitani. OP50 grown in LB was used as the standard feed for nematode cultivation according to Komura et al (Komura 2024). Nematodes were cultured on an peptone-free nematode growth media (pfNGM) with a lawn of OP50 at 20°C (Hoshino *et al*. 2008). For egg synchronization, L4-stage nematodes were collected, and two volumes of a lysis solution (1.25% NaClO, 0.2 M KOH) were added relative to the nematode volume. After vortexing, the mixture was washed three times with M9 buffer supplemented with 0.02% gelatin. The resulting egg suspension was dropped onto pfNGM plates seeded with OP50 and cultured at 20°C (Komura *et al*. 2013).

### Feeding Preference Assay

*C. elegans* individuals grown to the L4 stage by synchronized culture were collected from the petri dishes in approximately 2 mL of M9 buffer supplemented with 0.02% gelatin. The washed nematodes were placed at the center of a Nematode Growth Medium (NGM) agar plate, and 20 µL of test bacteria (OP50 and LAB) adjusted to an optical density at 660 nm (OD660) of 20 were spotted at equal intervals 2 cm away from the nematodes (Suppl. Figure 1)(Yu *et al*. 2015). The number of nematodes on each spot was counted after 3 hours at 20°C. The experiment was performed in duplicate with two independent trials, and the mean values were calculated. Similarly, the test bacteria adjusted to the same turbidity were heat-treated at 95°C. for 10 minutes using a heat block, and the feeding preference assay was conducted as described above.

### Determination of *C. elegans* Lifespan

On day 2 of synchronized culture, nematodes were harvested and washed with gelatin-supplemented M9 buffer. A mixture consisting of 10 mg of OP50 and FUdR (5-Fluoro-2’-deoxy-β-uridine) at a final concentration of 500 μg/mL in M9 buffer was inoculated onto pfNGM plates. On day 3 of synchronization, 50 of the FUdR-treated nematodes were transferred via a pick to pfNGM plates, where a mixture of FUdR and 10 mg of each bacterial test strain was applied directly to them. Survival was monitored every 3 to 4 days using a microscope. Nematodes were scored as alive if they showed spontaneous motility or responded to a physical touch stimulus with a pick. During each survival count, nematodes were transferred to fresh media containing the respective test strains as food, and this process was performed in triplicate for each strain. To feed the nematodes with heat-killed bacteria, a 100 μL mixture containing OP50 (10 mg/100 μL) and each heat-treated *L. lactis* strain (10 mg/100 μL) at a 1:1 ratio was prepared. After adding the prescribed volumes of FUdR solution and M9 buffer, this final mixture was introduced onto the pfNGM plates already seeded with the nematodes. Kaplan-Meier survival curves and statistical analyses were performed using the OASIS 2 (Online Application for Survival Analysis 2) software(Han *et al*. 2016). Statistical significance of differences in lifespan between groups was evaluated using the log-rank test with Bonferroni correction for multiple comparisons. Each experiment was replicated three or four times.

### Feeding assay of LAB I

Synchronized three-day-old adult *C. elegans* nematodes were harvested from NGM plates using 2 mL of M9 buffer supplemented with 0.02% gelatin and then washed three times with the same buffer. Each bacterial strain was suspended in M9 buffer to an optical density at 660 nm (OD_660_) of 10. For the 10- and 60-minute feeding assays, 100 μL of the bacterial suspension was spotted onto NGM plates; for the 300-minute assay, 300 μL was spotted. The feeding assay was initiated by transferring 30–40 mg of nematodes to each plate. At 10, 60, and 300 minutes after the onset of feeding, the nematodes were harvested with gelatin-supplemented M9 buffer, washed three times with the same buffer, and stored at −80°C.

### Feeding assay of LAB II

Nematodes were processed as described in *Feeding Assay I*, except that each bacterial strain was suspended to a final concentration of 10 mg/mL, and the feeding duration was fixed at 60 minutes. The treated nematodes were harvested, washed three times with gelatin-supplemented M9 buffer, and stored at −80°C.

### Isolation of total RNA

To each frozen aliquot of nematodes (in 1.5 mL microcentrifuge tubes), 350 µL of Buffer RA1 (NucleoSpin RNA extraction kit; Takara Bio, Shiga, Japan) and 3.5 µL of 2 M dithiothreitol (DTT) were added. An equal volume of zirconia beads (relative to the pellet volume of the nematodes) was added to the tube. The mixture was vortexed and frozen at −80°C for 10 minutes; this freeze-vortex cycle was performed three times. Total RNA was extracted using the NucleoSpin RNA extraction kit according to the manufacturer’s instructions. The concentration of the recovered RNA was determined using the Qubit 4 Fluorometer with the Qubit RNA BR Assay Kit (Thermo Fisher Scientific, Tokyo, Japan).

### Transcriptome analysis of nematodes fed with LAB

RNA sequencing was outsourced to Eurofins Genomics (Tokyo, Japan). and 2 × 150 bp paired-end sequencing was performed using the Illumina NovaSeq platform. From the obtained raw data, reads with a quality score of 15 or less for more than 40% of the read length, or those with a read length of 100 bp or less, were filtered out using fastp. Subsequently, the filtered reads were mapped to the *C. elegans* Bristol N2 reference genome (acc. no. GCF_000002985.6) using Bowtie2 (Langmead and Salzberg 2012) Based on the mapping results, the depth at each genomic position was calculated using Samtools (Li *et al*. 2009), and the Fragments Per Kilobase of exon per Million mapped fragments (FPKM) for each open reading frame (ORF) was determined (Danecek *et al*. 2021). To clarify the relative expression relationships between samples, principal component analysis (PCA) was performed on the FPKM values of each ORF using Scikit-learn (Pedregosa *et al*. 2011). Furthermore, differentially expressed gene (DEG) analysis was conducted using DESeq2 based on the FPKM data (Love *et al*. 2014). For functional characterization, re-annotation of each ORF was performed with eggNOG-mapper, and the ORFs were classified into Clusters of Orthologous Groups (COG) categories (Huerta-Cepas *et al*. 2017).

### RT-qPCR

Complementary DNA (cDNA) was synthesized using the QuantiTect Reverse Transcription Kit (Qiagen, Hilden, Germany). Quantitative PCR (qPCR) was performed using the Thermal Cycler Dice (TaKaRa, Shiga, Japan) with Thunderbird SYBR qPCR Mix (Toyobo, Osaka, Japan). The amplification program consisted of an initial denaturation at 95°C for 30 s, followed by 40 cycles of 95°C for 5 s and 60°C for 30 s. The following primer pairs were used for the reactions:

tuf: 5’-TGAAGAATTGATGGAACTCG-3’ (forward) / 5’-CATTGTGGTTCACCGTTC-3’ (reverse)

Arc Set 1: arcA43 5’-AAATCAGTCCTCCTCCACCG-3’ (forward) / arcA336 5’-TGAAGTCAAATATTCAGTCAAACCA-3’ (reverse)

Arc Set 2: arcA63 5’-CCCAGGTGCGGAAGTTGAA-3’ (forward) / arcA360 5’-TTCGACCATATCTTTTGTTGACATT-3’ (reverse)

Arc Set 3: arcA377 5’-CTATGCTGGCGTTCGTAAAAATGAA-3’ (forward) / arcA594 5’-ACGTGGGTGATTAGCCATAACA-3’ (reverse)

### Metabolome analysis of nematodes fed with LAB

Approximately 20-40 mg of frozen tissue of nematodes was placed in a homogenization tube, along with zirconia beads (5 mmφ and 3 mmφ). Next, 1,500 µL of 50% acetonitrile/Milli-Q water containing internal standards (H3304-1002, Human Metabolome Technologies, Inc. (HMT), Tsuruoka, Yamagata, Japan) was added to the tube, after which the tissue was completely homogenized at 1,500 rpm, for 60 sec ×15 at 4°C using a beads shaker. The homogenate was then centrifuged at 2,300 ×*g*, 4°C for 5 min. Subsequently, 400 µL of the supernatant was centrifugally filtered through a 5-kDa cutoff filter (HMT) at 9,100 ×*g*, 4°C to remove macromolecules. The filtrate was evaporated to dryness under vacuum and reconstituted in 50 µL of Milli-Q water for metabolome analysis.

Metabolome analysis was conducted according to HMT’s *Basic Scan* package, using capillary electrophoresis time-of-flight mass spectrometry (CE-TOFMS) based on the methods described previously (Ohashi *et al*. 2008; Ooga *et al*. 2011). Briefly, CE-TOFMS analysis was carried out using an Agilent CE capillary electrophoresis system equipped with an Agilent 6230 time-of-flight mass spectrometer (Agilent Technologies, Inc., Santa Clara, CA, USA). The systems were controlled by Agilent MassHunter Workstation Data Acquisition (Agilent Technologies) and connected by a fused silica capillary (50 μm *i.d.* × 62 cm total length) with commercial electrophoresis buffer (H3301-1001 and I3302-1023 for cation and anion analyses, respectively, HMT) as the electrolyte. The spectrometer was scanned from *m/z* 50 to 1,000 and peaks were extracted using MasterHands, automatic integration software (Keio University, Tsuruoka, Yamagata, Japan) in order to obtain peak information including *m/z*, peak area, and migration time (MT) (Sugimoto *et al*. 2010). Signal peaks corresponding to isotopomers, adduct ions, and other product ions of known metabolites were excluded, and the remaining peaks were annotated according to HMT’s metabolite database based on their *m*/*z* values and MTs. Areas of the annotated peaks were then normalized to internal standards and sample amount in order to obtain relative levels of each metabolite. Primary 110 metabolites were absolutely quantified based on one-point calibrations using their respective standard compounds. Hierarchical cluster analysis (HCA) and principal component analysis (PCA) (Yamamoto *et al*. 2014) were performed by HMT’s proprietary MATLAB and R programs, respectively. Detected metabolites were plotted on metabolic pathway maps using VANTED software(Junker *et al*. 2006).

### UPLC-MS analysis of citrulline, ornithine, and arginine

OP50 and LAB strains were harvested after cultivation, washed with PBS, and 10-mg aliquots of the bacterial pellets were stored at −80°C. To each frozen sample in a screw-cap tube, 0.5 g of zirconia beads and 100 µL of 50% (v/v) aqueous acetonitrile were added. Cell disruption was performed twice (4000 rpm for 30 s) using a cell homogenizer in a cold room. After disruption, the liquid phase was transferred to a new 1.5 mL microcentrifuge tube. To recover the remaining cell debris, 200 µL of 50% (v/v) acetonitrile was added to the beads, and the supernatant was pooled with the initial lysate; this washing step was performed twice. The combined lysate (approximately 500 µL) was centrifuged at 10,000 × *g* and 4°C for 5 min, and the resulting supernatant was filtered through a 0.22-µm membrane filter. A 100-µL aliquot of this bacterial extract was transferred to a 1.5 mL microcentrifuge tube, evaporated to dryness using a centrifugal evaporator (CVE-200D; EYELA, Tokyo, Japan), and stored at −20°C.

The dried residue was reconstituted in 20 µL of Milli-Q water, and a 10-µL aliquot (equivalent to 1/10 of the starting material) was subjected to derivatization. Amino acid derivatization was performed using the AccQ•Tag Ultra Derivatization Kit according to the manufacturer’s protocol (Waters, Milford, MA, USA). LC-MS/MS analysis was performed using an ACQUITY UPLC system coupled with a Xevo TQ-S tandem quadrupole mass spectrometer (Waters). Chromatographic separation was achieved on an ACQUITY UPLC column (2.1 × 100 mm, 1.7 *μ*m; Waters) with mobile phase A (prepared by diluting AccQ•Tag Ultra Eluent A with sterile water at a 1:20 ratio) and mobile phase B (AccQ•Tag Ultra Eluent B).

Arg, Cit, and ornithine (Orn) in samples were derivatized with 6-aminoquinolyl-N-hydroxysuccinimidyl carbamate using the AccQ-Taq Ultra Derivatization Kit according to the manufacturer’s protocol (Waters, Milford, MA, USA). Ultra-performance liquid chromatography-tandem mass spectrometry (UPLC-MS/MS) analysis was performed on an ACQUITY UPLC system coupled to a Xevo TQ-S mass spectrometer (Waters). Chromatographic separation was achieved using an AccQ-Tag Ultra Column (2.1 × 100 mm, 1.7 µm; Waters) maintained at 55°C. The sample injection volume was 1 µL, and the flow rate was set to 0.7 mL/min. The mobile phase consisted of a gradient mixture of AccQ-Tag Ultra Eluent A and B (Waters). The gradient program for eluent B was applied as follows: 0.1% at 0–0.54 min, 0.1–9.1% at 0.54–5.74 min, 9.1–21.2% at 5.74–7.74 min, 21.2–59.6% at 7.74–8.04 min, 59.6–90% at 8.04–8.05 min, 90% at 8.05–8.64 min, 90–0.1% at 8.64–8.73 min, and maintained at 0.1% until 9.50 min. Mass spectra were acquired in positive electrospray ionization (ESI+) mode, and the derivatives of arginine (Arg-Tag), citrulline (Cit-Tag), and ornithine (Orn-Tag) were detected using multiple reaction monitoring (MRM). The ion source parameters were optimized with a source temperature of 150°C, a capillary voltage of 0.5 kV, and a desolvation temperature of 500°C. The flow rates for the desolvation and cone gases were set at 1,000 L/h and 50 L/h, respectively. Compound-specific MRM transitions were monitored with the following parameters: m/z 345>171 for Arg-Tag (cone voltage: 30 V, collision energy: 20 eV), m/z 346>171 for Cit-Tag (cone voltage: 30 V, collision energy: 20 eV), and m/z for Orn-Tag (cone voltage: 20 V, collision energy: 20 eV).

## Results

### Feeding preferences

Under laboratory conditions, *C. elegans* N2 is maintained exclusively on OP50 as a food source. To evaluate the effects of LAB ingestion, the nematodes must consume the strains without displaying avoidance behavior. Thus, we assessed their preference toward various LAB strains (Suppl. Figure 1). Our results demonstrated that Lp1534 was equally preferred to OP50. In contrast, while significantly fewer nematodes migrated toward viable Lc114 compared to OP50, the heat-killed cells attracted a comparable number of nematodes to OP50. Similarly, preference assays using viable and heat-killed forms of Lc5805 and LcG50 revealed that the number of nematodes attracted to these test strains was significantly lower than to OP50 in all cases (Suppl. Figure 2).

### Lifespan assay

Lifespan assays were performed by feeding the nematodes with viable or heat-killed cells of the test strains in comparison with viable OP50. Log-rank test analysis based on Kaplan-Meier survival curves revealed that all LAB-supplemented groups, regardless of whether live or heat-killed cells were administered, exhibited a significant extension of lifespan compared to the OP50-fed control group (*p* < 0.001 for Lp1534, Lc5805, and LcG50; *p* < 0.01 for Lc114; Figure 1A–D).

**Figure 1.**
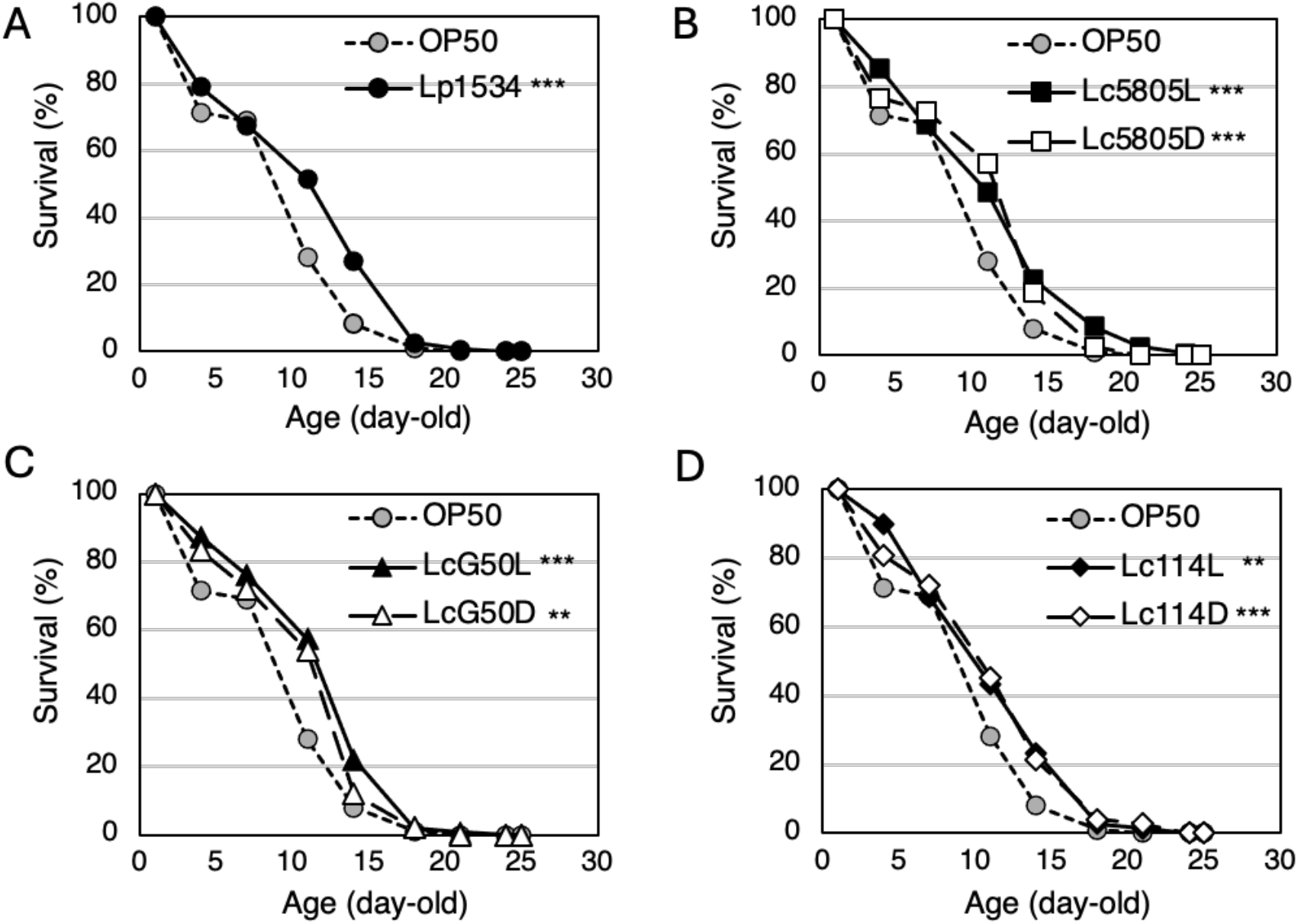
Effect of live or heat-killed LAB on the lifespan of *C. elegans*. (A–D) Survival curves of *C. elegans* fed (A) Lp1534, (B) live Lc5805 (Lc5805L) and heat-killed Lc5805 (Lc5805D), (C) live LcG50 (LcG50L) and heat-killed LcG50 (LcG50D), and (D) live Lc114 (Lc114L) and heat-killed Lc114 (Lc114D), shown in comparison with OP50-fed *C. elegans* as a control. Survival was scored at the indicated time points in the absence of FUdR treatment. The day of young adult stage was defined as day 0. Statistical significance was analyzed using the log-rank test based on Kaplan-Meier survival analysis with Bonferroni correction for multiple comparisons. \*\*\**p* < 0.001, \*\**p* < 0.01 vs. OP50 control. Data were pooled from four independent experiments for (A) (*n* = 200 worms per group) and three independent experiments for (B–D) (*n* = 150 worms per group). The same OP50 control data are shown in panels A–D for comparison.

Although overall survival curves between live and heat-killed groups did not show a statistically significant difference across the entire timeframe (*p* = 0.5622 for live vs. heat-killed Lc5805), distinct trends were observed in maximum survival times. While the viable OP50-fed group reached 100% mortality at 21 days, the viable Lc5805 (Lc5805L)-fed group exhibited the longest maximum survival, requiring 25 days for complete mortality (Figure 1B). In contrast, the heat-killed Lc5805 (Lc5805D)-fed group reached 100% mortality at 21 days, identical to the OP50 control. Similarly, for other *L. lactis* strains, viable LcG50 and Lc114 groups required 24 and 22 days for complete mortality, whereas their heat-killed counterparts required 21 and 23 days, respectively (Figure 1C, D).

### Transcriptome analysis of nematodes fed with LAB

To investigate the transcriptional changes, we performed transcriptome analysis using RNA purified from nematodes fed OP50, Lc5805 that exhibited a lifespan-extending effect, or Lp1534 that did not extend lifespan for 10 min, 1 h, or 5 h. Principal Coordinate Analysis (PCoA) revealed that the samples clustered clearly according to the ingested bacterial strain regardless of the feeding duration, indicating that the host gene expression profile was significantly altered depending on the bacterial species consumed (Figure 2).

**Figure 2.**
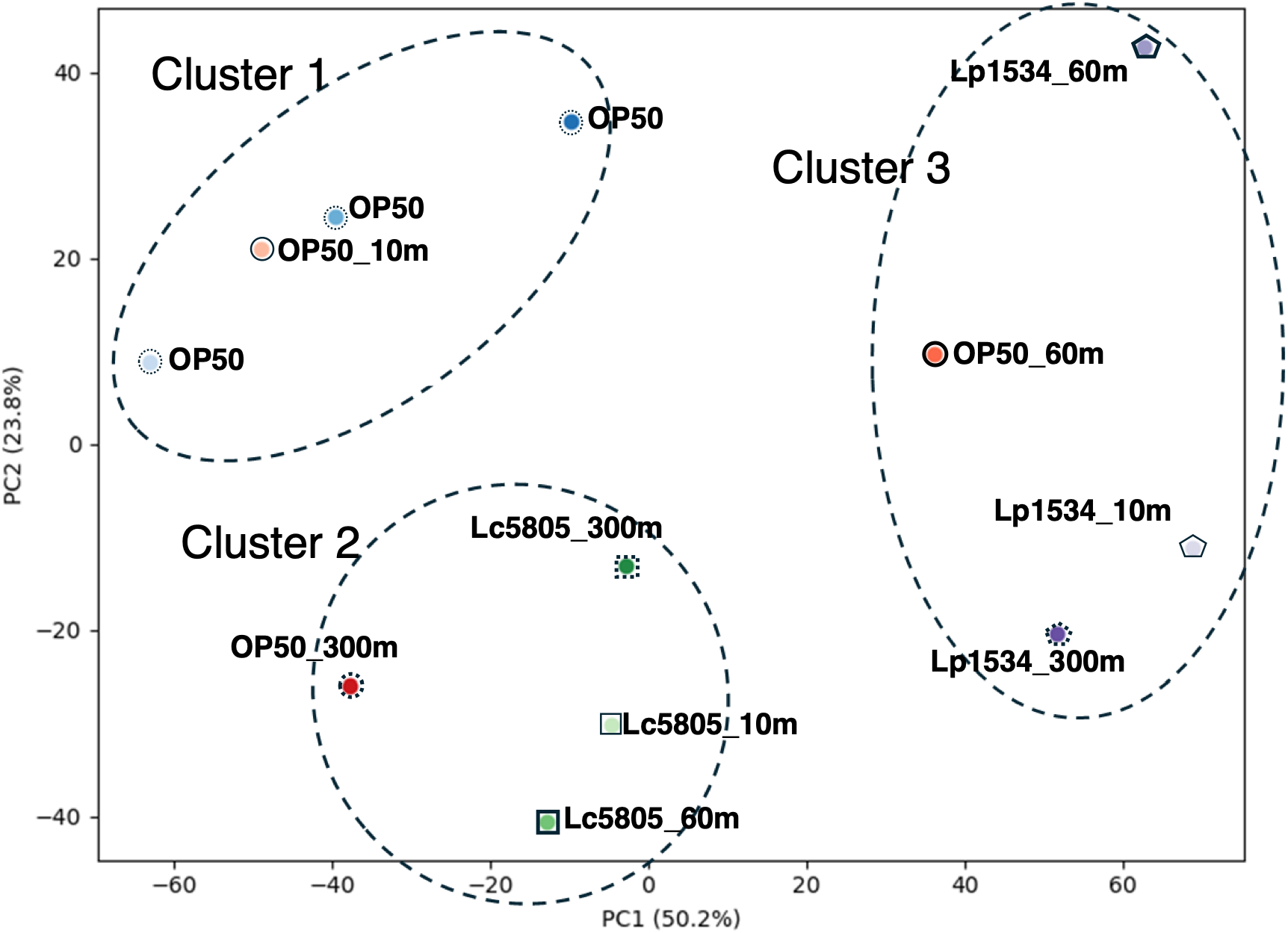
PCoA of gene expression profiles in *C. elegans* fed OP50, Lp1534, or Lc5805. PCoA clustering based on FPKM values successfully classified the samples into three distinct groups: cluster 1 (OP50-fed), cluster 2 (Lc5805-fed), and cluster 3 (Lp1534-fed).

Differentially expressed gene (DEG) analysis showed that the Lc5805-fed group had a larger number of upregulated genes compared to the OP50-fed group. Conversely, the Lp1534-fed group exhibited a larger number of downregulated genes than the OP50-fed group (Figure 3).

**Figure 3.**
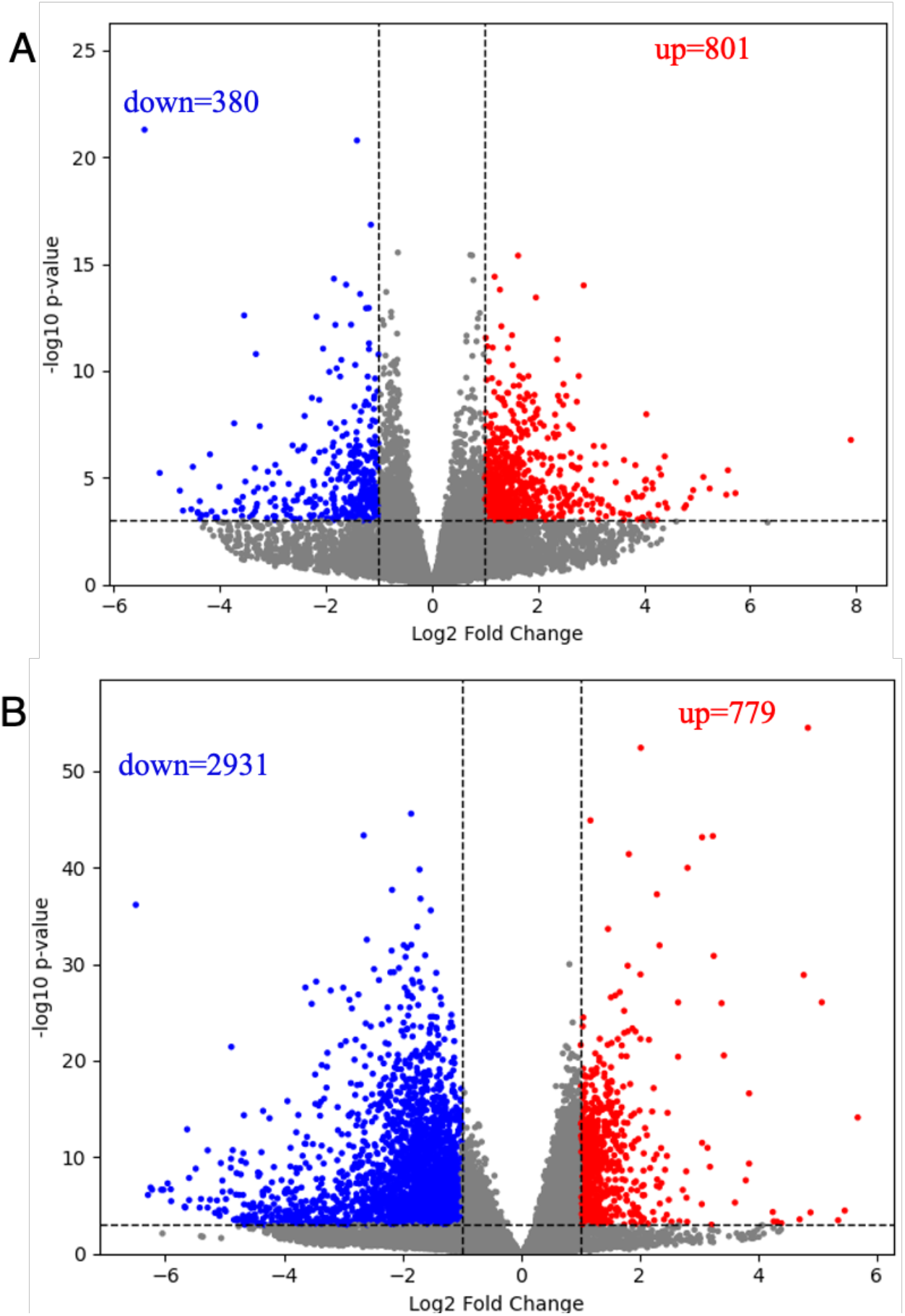
Volcano plots of differentially expressed genes in C. elegans fed OP50 vs. Lc5805 (A) and OP50 vs. Lp1534 (B). Differentially expressed gene (DEG) analysis was performed to identify genes exhibiting major changes in expression levels.

Furthermore, we categorized the DEGs using Clusters of Orthologous Groups (COG) analysis to compare functional variations. In the Lc5805-fed group, a substantial number of upregulated genes were classified into the following categories: “T: Signal transduction mechanisms,” “W: Extracellular structures,” “O: Post-translational modification, protein turnover, and chaperones,” and “P: Inorganic ion transport and metabolism” (Figure 4).

**Figure 4.**
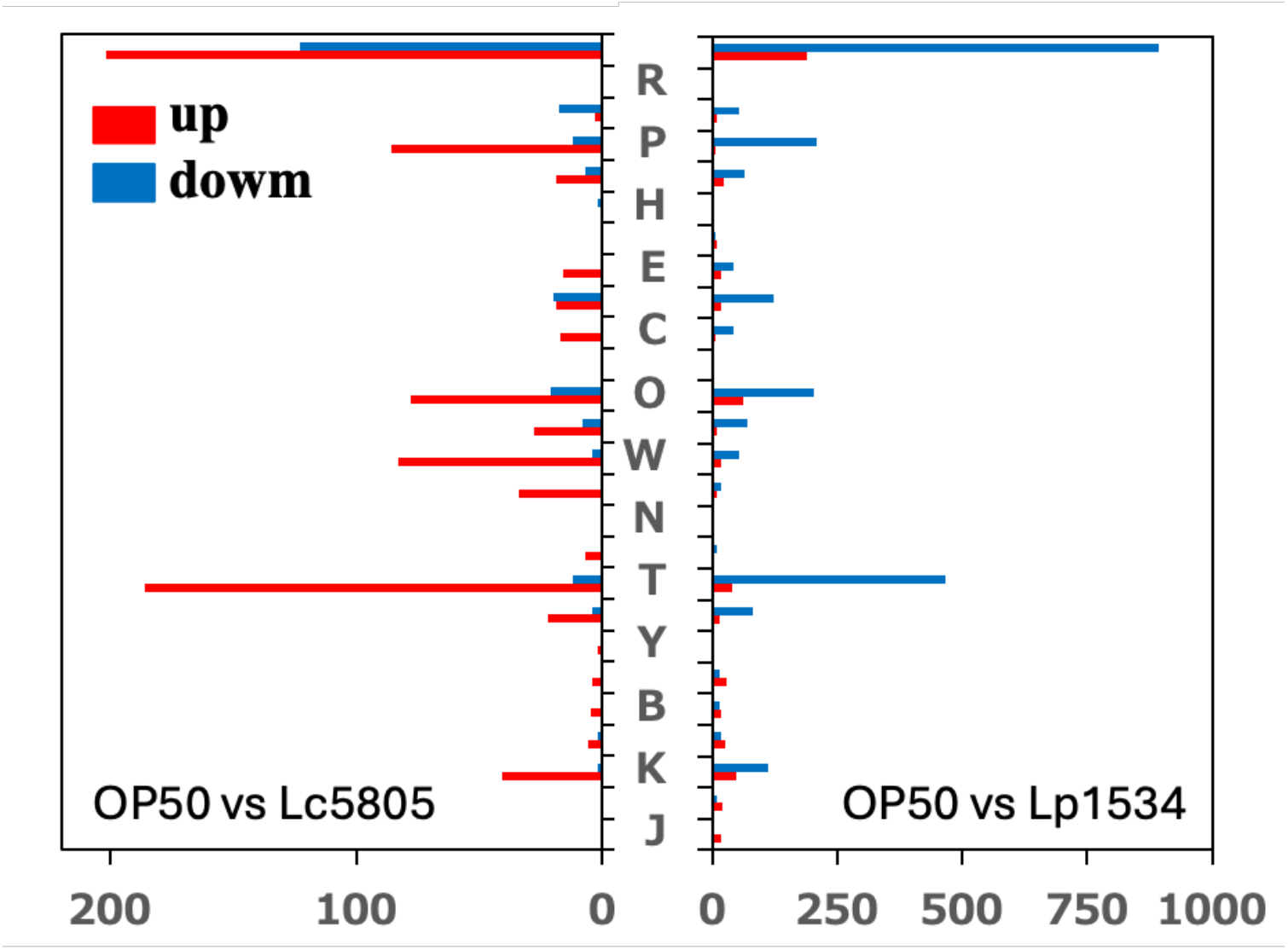
Differentially expressed genes categorized by COG functional classification. COG analysis of differentially expressed genes showed that, compared to OP50, Lc5805 showed increased expression in the CELLULAR PROCESSES AND SIGNALING (T, W, O) and METABOLISM (P) categories, while Lp1534 exhibited decreased expression in the same COG categories.

Specifically, within category W, which comprises 358 genes, 74 genes that were upregulated in the Lc5805-fed group remained unchanged or were downregulated in the Lp1534-fed group. All of these 74 genes are predicted to be involved in cuticle formation; among them, 58 were *col* genes, 3 were *bil* genes, and the remaining included *dpy-4*, *rol-1*, and genes of unknown function (Table 1).

**Table 1.** Upregulated genes in *C. elegans* fed Lc5805 relative to OP50 within COG category W (Extracellular structures).

| Category | Expression in Lp1534 (vs. OP50) | Gene symbol | Expression Pattern / Function in Cuticle |
|---|---|---|---|
| Collagens (*col* genes) | no | *col-3, 10, 12, 13, 14, 34, 40, 48, 51, 58, 65, 68, 74, 77, 81, 89, 97, 104, 105, 107, 109, 113, 115, 117, 125, 129, 130, 131, 139, 144, 145, 146, 157, 165, 167, 168, 169, 170, 172, 180* | Cuticle structural component (Adult-stage exoskeleton) |
|  | down | *col-2, 33, 36, 37, 38, 39, 44, 49, 50, 63, 71, 85, 110, 138, 141, 147, 175, 185* |  |
| Body Morphology | no | *dpy-1, dpy-4, ins-1, idpp-7, unc-122* | Cuticle integrity & body shape regulation |
| Other Cuticle Genes | no | *bil-6, sqt-2, ram-2* | Bilayer/cuticle structure |
|  | down | *bil-1*, *bil-2, rol-1* |  |

### Metabolome analysis of nematodes fed with LAB

To profile the metabolic changes, nematodes treated under the same conditions as those used for the transcriptome analysis (fed with OP50, Lc5805, or Lp1534 for 10 min, 1 h, or 5 h) were harvested and subjected to metabolome analysis using capillary electrophoresis-time-of-flight mass spectrometry (CE-TOFMS). Metabolomic analysis detected a wide range of metabolites across sample groups (Supplementary Table 1). Heatmap analysis visualized a distinct cluster of metabolites that specifically accumulated in the nematodes fed with Lc5805 for 60 min (Figure 5). Consistent with this, Principal Component Analysis (PCA) demonstrated that the 60-min Lc5805-fed group plotted at a noticeably distinct position compared to the other Lc5805-fed groups, the Lp1534-fed groups, and the OP50-fed control groups (Figure 6).

**Figure 5.**
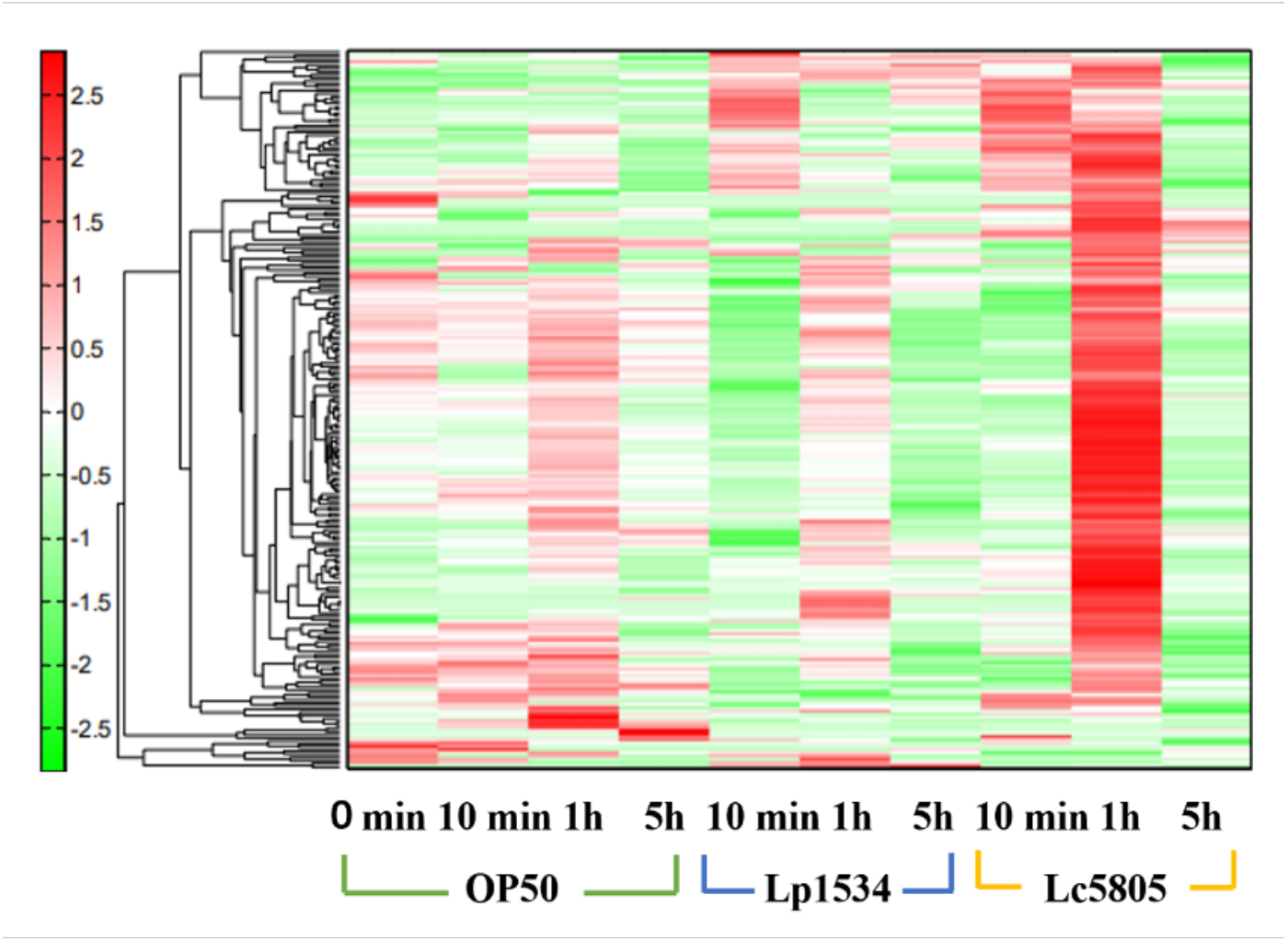
HeatMap of metabolites of *C. elegans* fed OP50, Lp1534, or Lc5805. Hierarchical Cluster Analysis (HCA) was performed for the peak of each test strain fed, and the distance between peaks is represented by the dendrogram in the figure.

**Figure 6.**
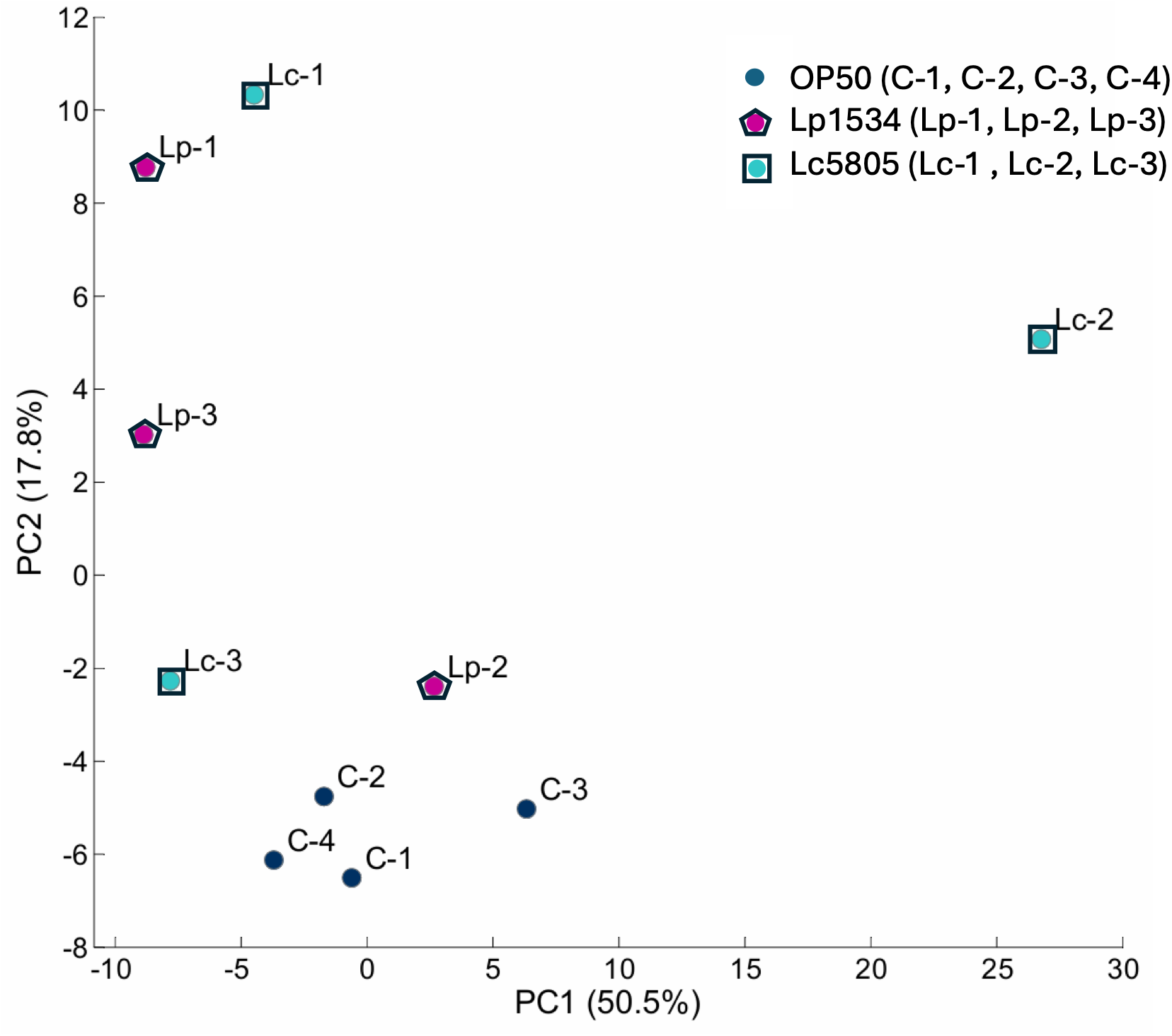
Principal component analysis (PCA) of metabolites in *C. elegans* fed OP50, Lp1534, or Lc5805. Two-dimensional PCA score plot based on CE-TOFMS metabolomic profiles of *C. elegans* fed OP50 (dark blue dots; C-1: 0 min, C-2: 10 min, C-3: 60 min, C-4: 300 min), Lp1534 (purple dots; Lp-1: 10 min, Lp-2: 60 min, Lp-3: 300 min), or Lc5805 (cyan dots; Lc-1: 10 min, Lc-2: 60 min, Lc-3: 300 min). The first and second principal components (PC1 and PC2) account for 50.5% and 17.8% of the total variance, respectively. Note the prominent separation of the 60-min Lc5805-fed group (Lc-2) along the PC1 axis.

The most notable metabolic change observed in this analysis was the remarkable accumulation of Cit in the Lc5805-fed groups. Because *C. elegans* lacks genes encoding most of the enzymes involved in the mammalian urea cycle, Cit is not endogenously synthesized in the host; indeed, its concentration remained at single-digit nmol/g levels in the OP50- and Lp1534-fed groups. In stark contrast, the Lc5805-fed groups exhibited a dramatic increase in Cit, exceeding 100 nmol/g at all time points (10, 60, and 300 min). Other metabolites that showed higher accumulation trends in the Lc5805-fed groups than in the OP50-fed group included hydroxyproline, asparagine, and aspartic acid. The accumulated levels of urea cycle-related amino acids—Arg, Orn, and Cit—along with other highly altered metabolites in each group are summarized in Table 2.

**Table 2.**
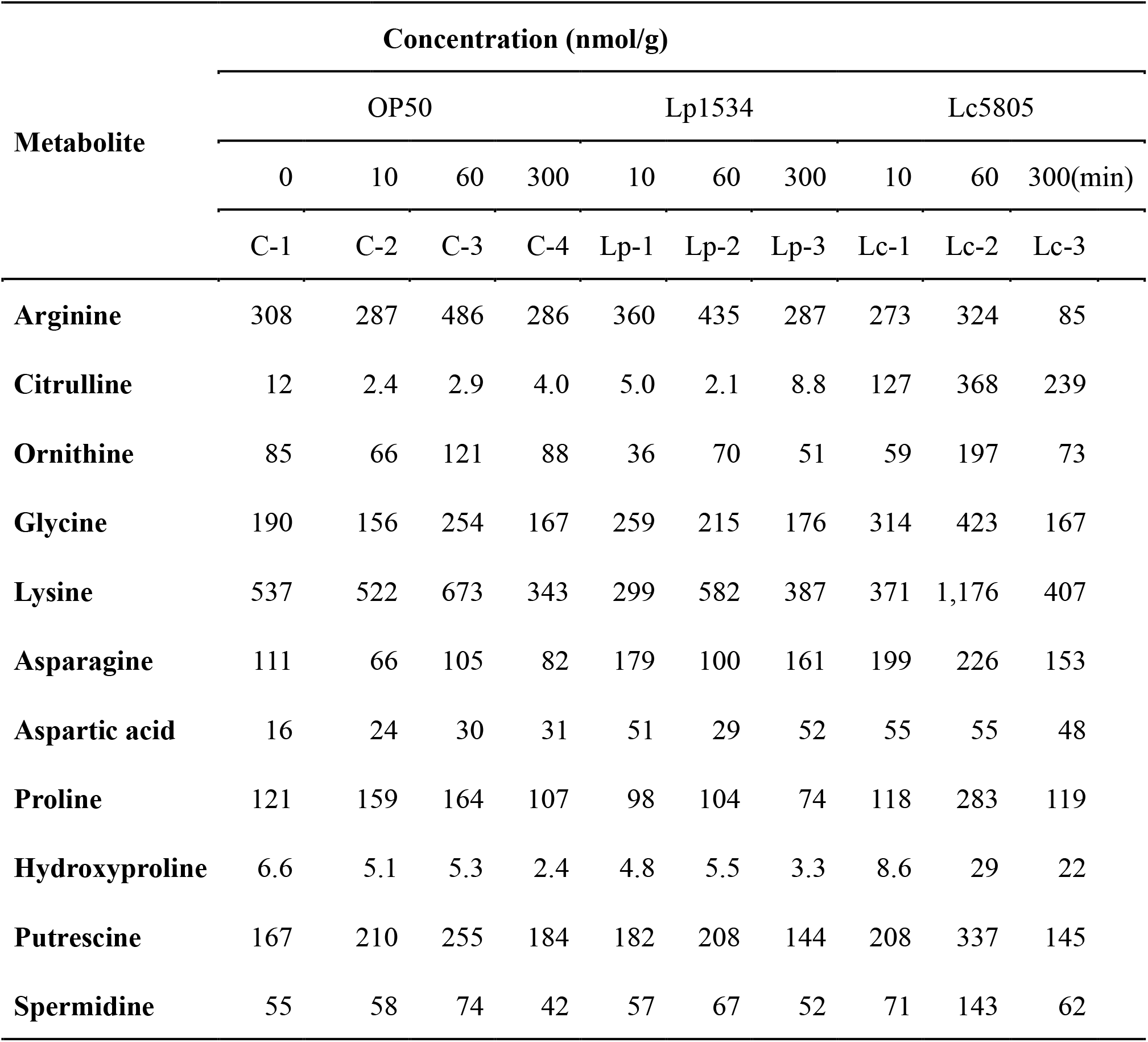
Concentrations of selected metabolites in *C. elegans* across different exposure times to lactic acid bacteria.

| **Metabolite** | **Concentration (nmol/g)** |  |  |  |  |  |  |  |  |  |
|---|---|---|---|---|---|---|---|---|---|---|
|  | OP50 |  |  |  | Lp1534 |  |  | Lc5805 |  |  |
|  | 0 | 10 | 60 | 300 | 10 | 60 | 300 | 10 | 60 | 300(min) |
|  | C-1 | C-2 | C-3 | C-4 | Lp-1 | Lp-2 | Lp-3 | Lc-1 | Lc-2 | Lc-3 |
| **Arginine** | 308 | 287 | 486 | 286 | 360 | 435 | 287 | 273 | 324 | 85 |
| **Citrulline** | 12 | 2.4 | 2.9 | 4.0 | 5.0 | 2.1 | 8.8 | 127 | 368 | 239 |
| **Ornithine** | 85 | 66 | 121 | 88 | 36 | 70 | 51 | 59 | 197 | 73 |
| **Glycine** | 190 | 156 | 254 | 167 | 259 | 215 | 176 | 314 | 423 | 167 |
| **Lysine** | 537 | 522 | 673 | 343 | 299 | 582 | 387 | 371 | 1,176 | 407 |
| **Asparagine** | 111 | 66 | 105 | 82 | 179 | 100 | 161 | 199 | 226 | 153 |
| **Aspartic acid** | 16 | 24 | 30 | 31 | 51 | 29 | 52 | 55 | 55 | 48 |
| **Proline** | 121 | 159 | 164 | 107 | 98 | 104 | 74 | 118 | 283 | 119 |
| **Hydroxyproline** | 6.6 | 5.1 | 5.3 | 2.4 | 4.8 | 5.5 | 3.3 | 8.6 | 29 | 22 |
| **Putrescine** | 167 | 210 | 255 | 184 | 182 | 208 | 144 | 208 | 337 | 145 |
| **Spermidine** | 55 | 58 | 74 | 42 | 57 | 67 | 52 | 71 | 143 | 62 |

### Kinetic analysis of urea cycle-related metabolites

Although metabolome analysis revealed a remarkable accumulation of Cit in the Lc5805-fed group, the underlying metabolic pathway remained unclear. To determine whether this Cit accumulation was directly derived from the ingested Lc5805 cells or synthesized within the nematode body via Lc5805-derived enzymes, we performed quantitative UPLC-MS analysis of urea cycle-related metabolites—Arg, Cit, and Orn—using both bacterial cells alone and nematodes fed with either live or heat-killed bacteria. To further investigate whether this Cit accumulation was a specific response to Lc5805, we expanded our bacterial panel. In addition to OP50, Lp1534, and Lc5805, we included two other *L. lactis* strains belonging to the same species as Lc5805: Lc114 and LcG50.

In the quantitative analysis of the bacterial cells alone, Arg was found to accumulate in OP50 and Lp1534, whereas Orn accumulated in Lc5805, LcG50, and Lc114. Notably, Cit was detected exclusively in Lc5805 cells (Figure 7A). Meanwhile, in nematodes fed with either live or heat-killed bacteria, a significantly higher accumulation of Cit was observed in the groups fed with live Lc5805 and live LcG50 (Figure 7B). In contrast, no Cit accumulation was detected in the groups fed with live OP50, live Lp1534, heat-killed Lp1534, or heat-killed Lc5805. For Lc114, Cit accumulation was observed in both the live- and heat-killed-fed groups. In the groups fed with live Lc5805 and live LcG50, all three metabolites (Arg, Cit, and Orn) were detected. However, in the groups fed with their heat-killed counterparts, only Arg was detected. This observation suggests that the conversion of Arg to Cit was inactive in the heat-killed groups, potentially explaining the decreased accumulation of Arg in the live-fed groups. Collectively, these results suggest that the Cit accumulated in the nematodes fed with Lc5805 is likely produced by the bacterial cells themselves and/or their intracellular enzymes.

**Figure 7.**
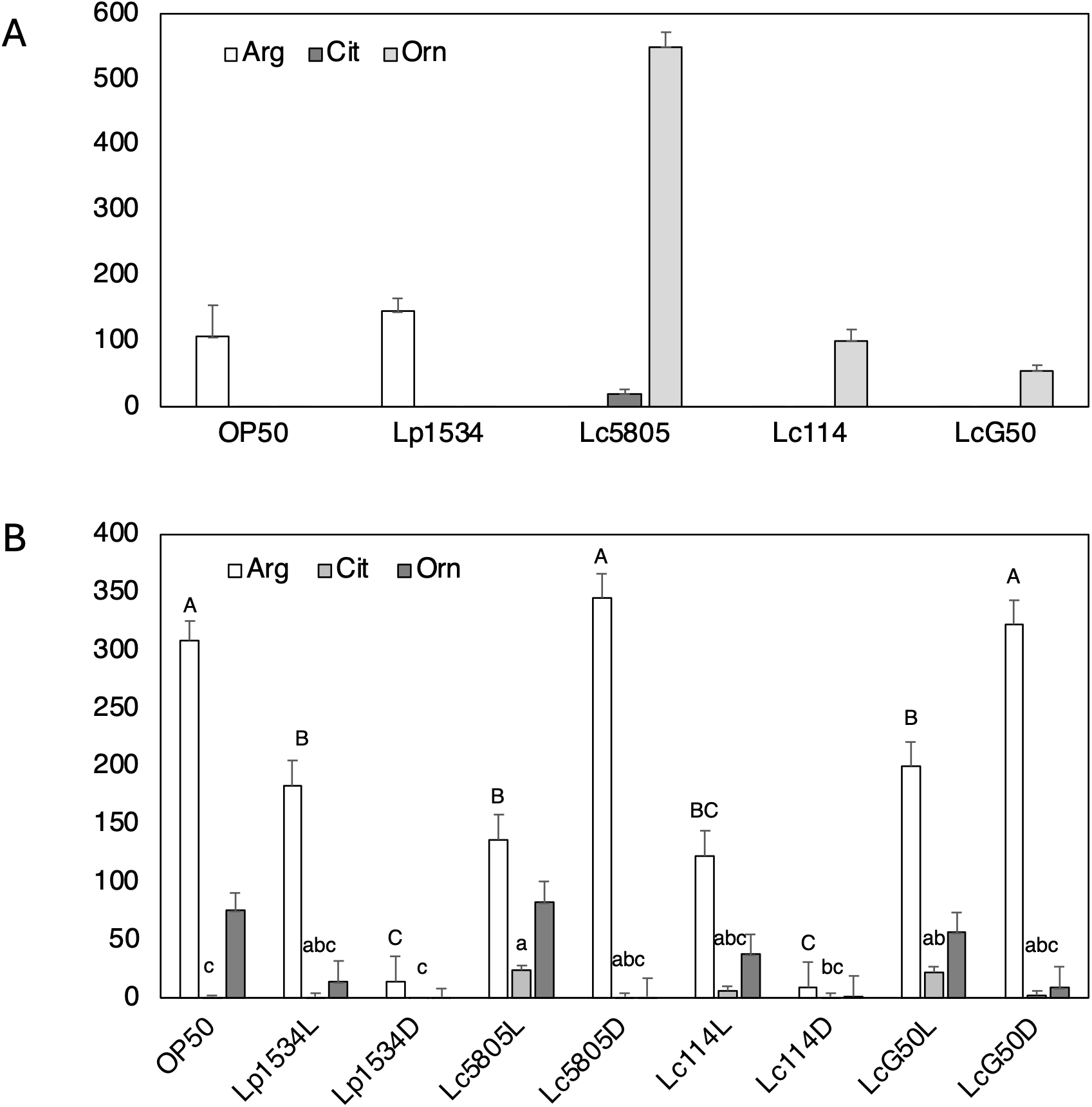
Quantification of Arg, Cit, and Orn in cell lysate of test strains (A) and in *C. elegans* extracts fed with each test strain (B). The “L” or “D” following the strain name indicates live or dead bacteria, respectively. Different uppercase letters indicate significant differences in Arg content (p < 0.05, Tukey’s HSD test). Different lowercase letters indicate significant differences in Cit content (p < 0.05, Tukey’s HSD test). No significant differences were found in Orn content (p < 0.05, Tukey’s HSD test).

Focusing on the *L. lactis arcA* gene, which encodes arginine deiminase—the enzyme responsible for the first step of the arginine deiminase pathway that hydrolyzes Arg into Cit and ammonia (Larsen *et al*. 2004)—we performed RT-qPCR using total RNA isolated from the respective nematodes. The *L. lactis tuf* gene was used as an internal reference (housekeeping gene), and the expression of the *arcA* gene was measured using three distinct primer sets. When calculating the Δ*C_t_* values for each sample, no values were obtained for some samples in the OP50-fed control group. Conversely, negative Δ*C_t_* values were consistently calculated for all Lc5805-fed samples (Table 3), indicating very high expression of the *arcA* gene derived from Lc5805 within the nematode. Furthermore, the *C_t_* values for both *arcA* and *tuf* were lower in the 10-min feeding group than in the 300-min feeding group, suggesting that the expression level of the *L. lactis arcA* gene within the nematode body decreased over time.

**Table 3.** qPCR of *L. lactis arcA* in *C. elegans* fed Lc5805.

| Fed strain and time | Ct | Primer set | Δ Ct |
|---|---|---|---|
| Lc5805<br>10 min | 25.3 | 5' arc 43-3' arc 336 | -3.66 |
|  | 24.95 | 5' arc 63-3' arc 360 | -4.01 |
|  | 24.4 | 5' arc 367-3' arc 594 | -4.56 |
|  | 28.96 | tuf | 0 |
| Lc5805<br>300 min | 31.76 | 5' arc 43-3' arc 336 | -1.29 |
|  | 30.62 | 5' arc 63-3' arc 360 | -2.43 |
|  | 29.14 | 5' arc 367-3' arc 594 | -3.91 |
|  | 33.05 | tuf | 0 |

## Discussion

In this study, we evaluated the effects of daily ingestion of LAB on healthspan and elucidated its underlying molecular mechanisms using *C. elegans* as an alternative model system to mammalian rodents.

Because laboratory *C. elegans* are conventionally maintained solely on *E. coli* OP50, we first evaluated the dietary preference of *C. elegans* toward LAB strains and selected Lp1534, which exhibited food preference comparable to that for OP50. Subsequent lifespan assays using Lp1534 and Lc5805 revealed that while both LAB strains significantly extended mean lifespan, Lc5805 uniquely conferred a significantly longer maximum lifespan. To unravel the distinct physiological responses, we conducted time-course transcriptomic and metabolomic analyses on nematodes fed OP50, Lp1534, or Lc5805.

Transcriptomic profiling revealed that samples clustered primarily by the ingested bacterial strain rather than the feeding duration, indicating that the bacterial diet exerts a dominant influence on the host transcriptional landscape. Detailed COG functional classification demonstrated that among 358 genes in Category W (Extracellular structures), 74 genes specifically upregulated in the Lc5805 group (which remained unchanged or downregulated in the Lp1534 group) were predominantly involved in cuticle formation. In *C. elegans*, the cuticle serves as an exoskeleton and physical barrier, and the induction of collagen (*col*) genes is strongly linked to longevity. Intriguingly, despite the marked upregulation of numerous collagen genes upon Lc5805 consumption, host endogenous proline biosynthetic enzymes (*oatr-1*, *pycr-1*) and classical longevity/stress-response markers (*sod-3*, *gst-4*) showed no significant induction.

Metabolomic profiling provided the critical key to resolving this paradox. The most striking metabolomic feature was the specific accumulation of Cit and Orn exclusively in nematodes fed live Lc5805. Although single-agent L-Cit supplementation has been reported to extend *C. elegans* healthspan (Rajcic 2023), its physiological origin and mechanism remained elusive. Here, the detection of *L. lactis* arginine deiminase (*arcA*) mRNA within the nematode intestine strongly indicates that the accumulated Cit and Orn were not synthesized endogenously by the host but were directly provided by active Lc5805 through a combination of bacterial “metabolites + enzymes.” In *L. lactis*, *arcA* forms a conserved gene cluster (the arginine deiminase (ADI) operon) together with *arcB*, which encodes ornithine transcarbamylase (OTC) responsible for converting Cit to Orn. Because genes within this operon are co-transcribed, high expression of *arcA* strongly implies concurrent transcription of *arcB*, providing a rational mechanism for the conversion of Cit to Orn within the gut. The supplied Orn likely served as an exogenous precursor for collagen-constituent amino acids (e.g., proline)(Albaugh 2017; Barbul 2008), thereby supporting efficient cuticle maintenance without demanding host biosynthetic energy.

While heat-treated *L. lactis* (Lc5805D) confers a baseline lifespan extension comparable to live cells in terms of mean lifespan, likely mediated through heat-stable cell wall components and nutritional factors, it failed to extend the maximum lifespan beyond that of the control group (100% mortality at 21 days). In contrast, continuous ingestion of live *L. lactis* (Lc5805L) extended the maximum lifespan to 25 days. This distinction suggests that while structural bacterial components can enhance overall average healthspan, the active operation of the intestinal ADI pathway and continuous metabolite provision by live bacteria are indispensable for sustaining cuticle integrity and supporting late-life survival during advanced age.

Nitric oxide (NO) is a vital signaling molecule in multicellular organisms, typically synthesized from Arg by nitric oxide synthase (NOS). Lacking both NOS and a functional urea cycle, *C. elegans* reportedly “hijacks” bacterial NO generated by NOS-bearing *Bacillus subtilis* (*nosA*) in its natural habitat to enhance lifespan and stress tolerance(Gusarov 2013). However, *L. lactis* lacks a NOS gene. Thus, *L. lactis* promotes longevity through a novel, NO-independent mechanism distinct from that of *Bacillus*.

Collectively, these examples involving *Bacillus* NOS and *L. lactis* ADI suggest a broader ecological paradigm: in their natural habitat, *C. elegans* has evolutionarily adapted to utilize not only bacterial nutrients but also active bacterial enzymes to support its physiological functions. Because standard laboratory *C. elegans* is maintained exclusively on *E. coli* OP50, providing probiotic bacteria equipped with functional metabolic enzymes likely unlocks this evolutionary capacity, thereby conferring marked healthspan benefits.

In conclusion, these findings demonstrate that Lc5805-mediated longevity does not rely on host metabolic reprogramming, but rather on “Metabolic Exploitation,” wherein the host directly utilizes bacterially derived metabolites and enzymes. The finding that Cit accumulation was completely absent in nematodes fed OP50, the ADI-lacking strain Lp1534, or heat-treated Lc5805 confirms that this exogenous metabolic provisioning system, the operation of the ADI pathway, specifically requires active bacterial enzymes from live Lc5805 in the gut. By exploiting external amino acid and enzymatic resources, the host avoids the energetic costs of overactivating endogenous stress responses (e.g., *sod-3*, *gst-4*) while maintaining the structural integrity of the cuticle and supporting proteostasis. This study unveils a novel paradigm in host–microbe metabolic interactions supporting host healthspan and longevity.

## Supplementary material

Supplementary Table 1. Quantitative analysis of target metabolites

Supplementary Figure 1. Results of feeding preferences assay of live and heat-killed various LAB strains.

Supplementary Figure 2. Results of feeding preferences assay of live and heat-killed Lp5805 and Lc G50.

## Data availability

The RNA sequence data were deposited in the GenBank/EMBL/DDBJ database under bioproject number PRJDB45693. All target metabolite quantification data generated in this study are provided in Supplementary Table 1.

## Author Contribution

A.I., D.K. and H.M. designed and conducted the experiments, and analyzed the data. T.K. assisted with discussions. C.S. wrote the manuscript. C.S. supervised the project and contributed to the study design. All authors read and approved the final manuscript.

## Acknowledgments

We thank Prof. Shohei Mitani (Tokyo Women’s Medical University) for generously providing *Caenorhabditis elegans* and *Escherichia coli* OP50, and Dr. Tomomi Komura (University of Hyogo) for valuable guidance on *C. elegans* experimental procedures.

