## Supplemental figures and Table for "Metabolic exploitation of the bacterial arginine deiminase pathway extends *Caenorhabditis elegans* healthspan"

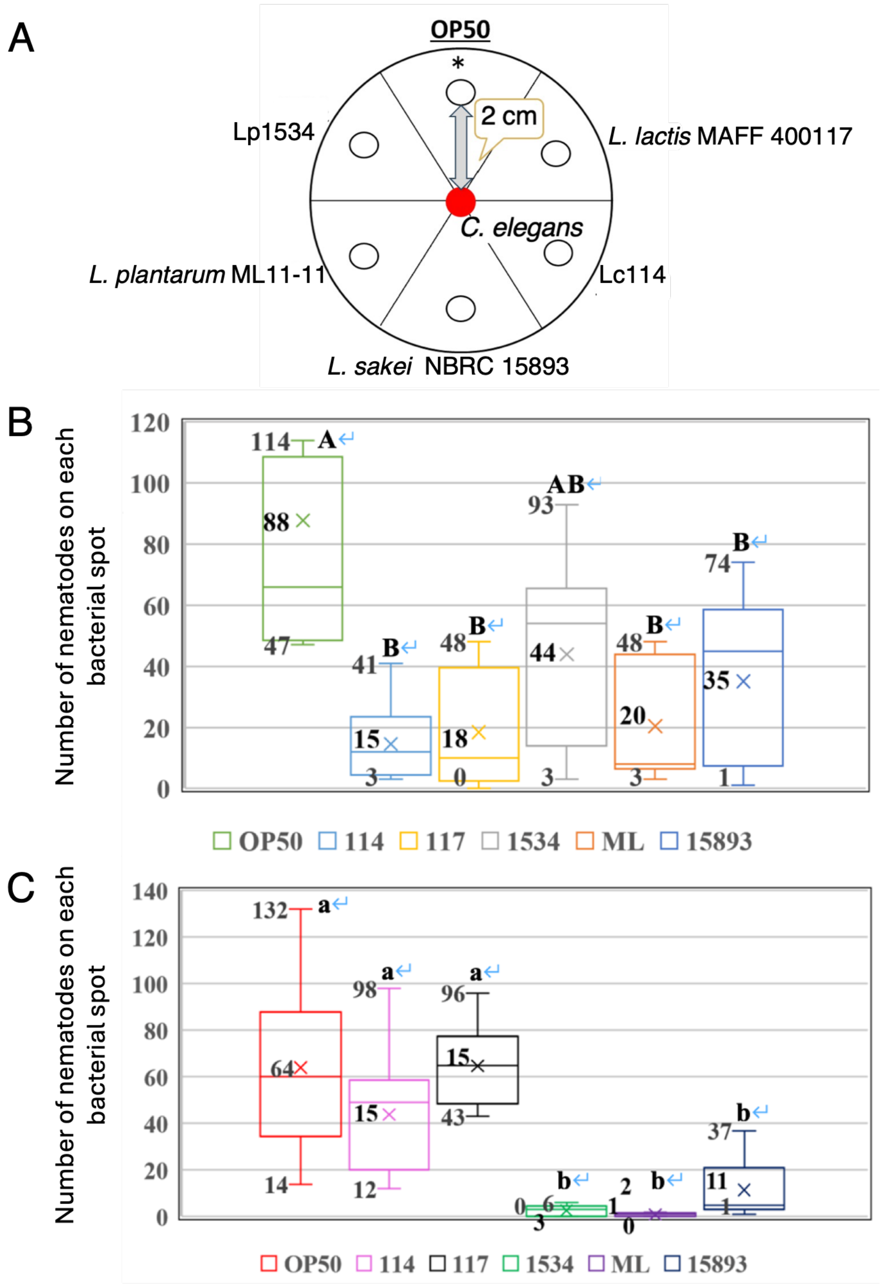


Supplementary Figure 1

(A) Schematic illustration of the food preference assay plate. Circles represent spots of different strains as indicated. (B, C) Number of nematodes on each bacterial spot containing (B) live or (C) heat-killed LAB strains. In the box plots, horizontal lines and crosses indicate medians and means, respectively; boxes represent the interquartile ranges, and whiskers indicate the minimum and maximum values. Different letters (uppercase in B, lowercase in C) indicate statistically significant differences among bacterial spots ($p < 0.05$, Tukey's HSD test).

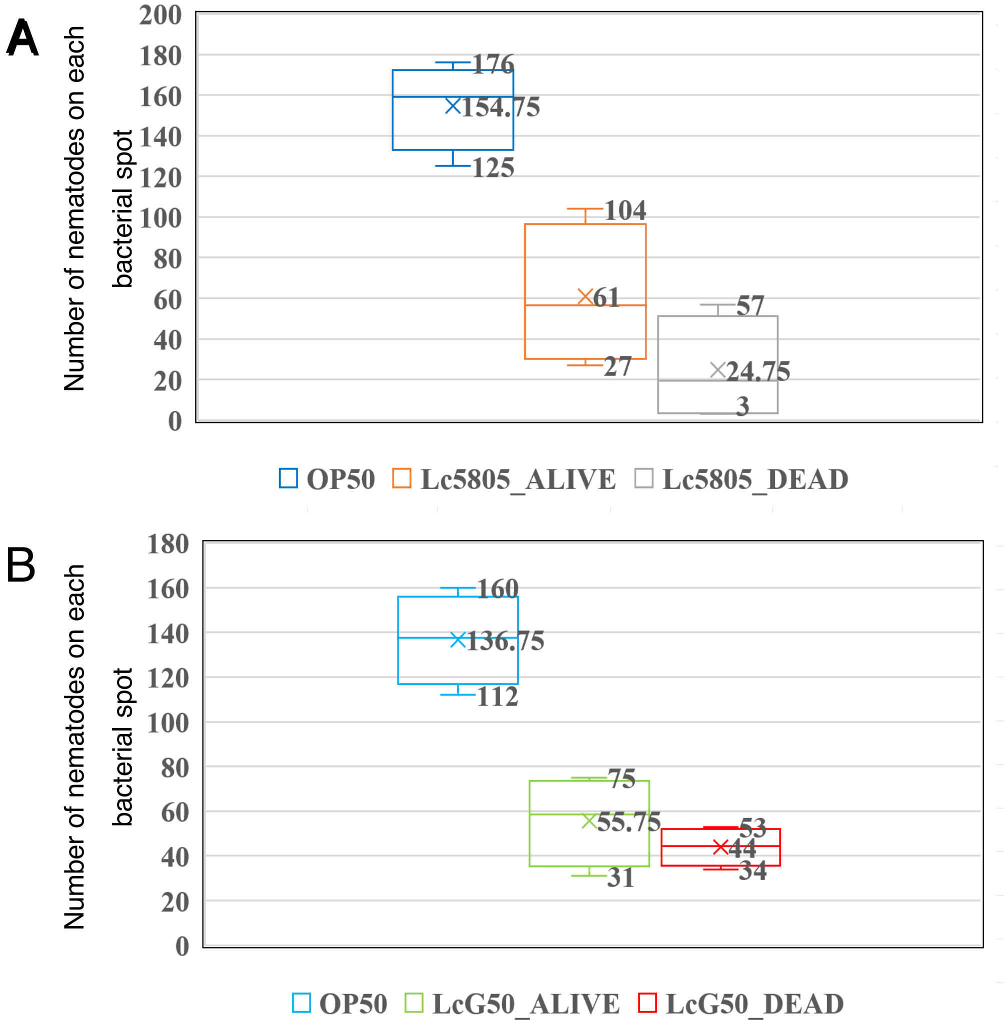


Supplementary Figure 2

(A) Number of nematodes on OP50, live Lc5805 and heat-killer Lc 5805. (B) Number of nematodes on OP50, live LcG50 and heat-killer LcG50. In the box plots, horizontal lines and crosses indicate medians and means, respectively; boxes represent the interquartile ranges, and whiskers indicate the minimum and maximum values.

### Supplementary Table S1. Quantitative analysis of target metabolites

| ID | Metabolite | PubChem CID | HMDB ID | Concentration (nmol/g) | | | | | | | | | |
|---|---|---|---|---|---|---|---|---|---|---|---|---|---|
| | | | | Control | | | | Lp1534 | | | Lc5805 | | |
| | | | | 0 min | 10 min | 60 min | 300 min | 10 min | 60 min | 300 min | 10 min | 60 min | 300 min |
| | | | | C-1 | C-2 | C-3 | C-4 | Lp-1 | Lp-2 | Lp-3 | Lc-1 | Lc-2 | Lc-3 |
| A_0080 | Acetyl CoA_divalent | 444493 | | 2.5 | 3.6 | 3.3 | 2.2 | 1.8 | 2.2 | 0.9 | 1.4 | 4.9 | 1.6 |
| A_0043 | *cis*-Aconitic acid | 643757 | HMDB0000072 | 3.5 | N.D. | 3.2 | 2.2 | N.D. | 2.9 | 2.7 | 2.5 | 5.1 | 2.8 |
| C_0048 | Adenine | 190 | HMDB0000034 | N.D. | N.D. | N.D. | N.D. | N.D. | N.D. | N.D. | N.D. | N.D. | N.D. |
| C_0133 | Adenosine | 60961 | HMDB0000050 | 0.4 | 0.4 | 0.4 | 0.3 | N.D. | 0.3 | 0.2 | 0.5 | 0.5 | N.D. |
| A_0086 | ADP | 6022 | HMDB0001341 | 116 | 107 | 133 | 94 | 79 | 114 | 86 | 100 | 202 | 84 |
| C_0007 | Ala | 602 | HMDB0000161 | 1,652 | 2,391 | 2,345 | 1,269 | 2,462 | 1,878 | 1,380 | 2,835 | 3,677 | 1,141 |
| A_0072 | AMP | 6083 | HMDB0000045 | 60 | 54 | 60 | 46 | 31 | 57 | 40 | 52 | 100 | 40 |
| C_0050 | Anthranilic acid | 227 | HMDB0001123 | 1.0 | 1.1 | 0.8 | 0.6 | N.D. | 1.1 | 0.8 | 0.4 | 1.2 | 0.4 |
| C_0082 | Arg | 6322 | HMDB0000517 | 308 | 287 | 486 | 286 | 360 | 435 | 287 | 273 | 324 | 85 |
| C_0044 | Asn | 236 | HMDB0000168 | 111 | 66 | 105 | 82 | 179 | 100 | 161 | 199 | 226 | 153 |
| C_0047 | Asp | 424 | HMDB0000191 | 16 | 24 | 30 | 31 | 51 | 29 | 52 | 55 | 55 | 48 |
| A_0095 | ATP | 5957 | HMDB0000538 | 273 | 280 | 376 | 253 | 254 | 276 | 197 | 225 | 557 | 196 |
| C_0025 | Betaine | 247 | HMDB0000043 | 202 | 160 | 181 | 125 | 308 | 180 | 174 | 357 | 374 | 165 |
| C_0029 | Betaine aldehyde_+$H_2O$ | 249 | | N.D. | N.D. | N.D. | N.D. | N.D. | N.D. | N.D. | N.D. | N.D. | N.D. |
| A_0069 | cAMP | 6076 | HMDB0000058 | N.D. | N.D. | N.D. | N.D. | N.D. | N.D. | N.D. | N.D. | N.D. | N.D. |
| C_0110 | Carnosine | 439224 | HMDB0000033 | N.D. | N.D. | N.D. | N.D. | N.D. | N.D. | N.D. | N.D. | N.D. | N.D. |
| A_0078 | CDP | 6132 | HMDB0001546 | 6.4 | 5.7 | 7.1 | 6.1 | 4.5 | 7.9 | 6.1 | 5.3 | 12 | 5.6 |
| A_0071 | cGMP | 24316 | HMDB0001314 | N.D. | N.D. | N.D. | N.D. | N.D. | N.D. | N.D. | N.D. | N.D. | N.D. |
| C_0016 | Choline | 305 | HMDB0000097 | 11 | 7.3 | 8.3 | 5.8 | 15 | 14 | 18 | 12 | 18 | 4.8 |
| A_0049 | Citric acid | 311 | HMDB0000094 | 148 | 126 | 184 | 127 | 172 | 146 | 155 | 189 | 251 | 129 |
| C_0083 | Citrulline | 9750 | HMDB0000904 | 12 | 2.4 | 2.9 | 4.0 | 5.0 | 2.1 | 8.8 | 127 | 368 | 239 |
| A_0067 | CMP | 6131 | HMDB0000095 | 5.3 | 4.3 | 5.1 | 3.9 | N.D. | 5.7 | 3.7 | 5.5 | 10 | 2.9 |
| A_0075 | CoA_divalent | 87642 | | 1.9 | N.D. | 2.3 | 2.2 | N.D. | 2.9 | 1.3 | 1.8 | 4.5 | 2.0 |
| C_0041 | Creatine | 586 | HMDB0000064 | N.D. | N.D. | N.D. | N.D. | N.D. | N.D. | N.D. | N.D. | N.D. | N.D. |
| C_0022 | Creatinine | 588 | HMDB0000562 | N.D. | N.D. | N.D. | N.D. | N.D. | N.D. | N.D. | N.D. | N.D. | N.D. |
| A_0091 | CTP | 6176 | HMDB0000082 | 16 | 17 | 25 | 19 | 15 | 18 | 13 | 13 | 38 | 13 |
| C_0030 | Cys | 594 | HMDB0000574 | N.D. | N.D. | N.D. | N.D. | N.D. | N.D. | N.D. | N.D. | N.D. | N.D. |
| C_0117 | Cytidine | 6175 | HMDB0000089 | 0.5 | 0.6 | 1.0 | 0.8 | N.D. | 1.1 | 0.5 | 0.6 | 1.8 | 0.8 |
| C_0020 | Cytosine | 597 | HMDB0000630 | N.D. | N.D. | N.D. | N.D. | N.D. | 0.8 | 1.4 | N.D. | N.D. | N.D. |
| A_0039 | Dihydroxyacetone phosphate | 668 | HMDB0001473 | N.D. | N.D. | 6.3 | 10 | 9.2 | 15 | 15 | 10 | 18 | 9.1 |
| C_0013 | *N*,*N*-Dimethylglycine | 673 | HMDB0000092 | N.D. | N.D. | N.D. | N.D. | N.D. | N.D. | N.D. | N.D. | N.D. | N.D. |
| A_0094 | dATP | 15993 | HMDB0001532 | N.D. | N.D. | N.D. | N.D. | N.D. | N.D. | N.D. | N.D. | N.D. | N.D. |
| A_0089 | dCTP | 65091 | HMDB0000998 | N.D. | N.D. | N.D. | N.D. | N.D. | N.D. | N.D. | N.D. | N.D. | N.D. |
| A_0077 | dTDP | 164628 | HMDB0001274 | N.D. | N.D. | N.D. | N.D. | N.D. | N.D. | N.D. | N.D. | N.D. | N.D. |
| A_0066 | dTMP | 9700 | HMDB0001227 | N.D. | N.D. | N.D. | N.D. | N.D. | N.D. | N.D. | N.D. | N.D. | N.D. |
| A_0090 | dTTP | 64968 | HMDB0001342 | N.D. | N.D. | N.D. | N.D. | N.D. | N.D. | N.D. | N.D. | N.D. | N.D. |
| A_0053 | Erythrose 4-phosphate | 122357 | HMDB0001321 | N.D. | N.D. | N.D. | N.D. | N.D. | N.D. | N.D. | N.D. | N.D. | N.D. |
| A_0070 | Fructose 1,6-diphosphate | 172313 | HMDB0001058 | N.D. | N.D. | N.D. | N.D. | N.D. | 2.2 | N.D. | N.D. | 2.6 | N.D. |
| A_0061 | Fructose 6-phosphate | 603 | HMDB0000124 | 12 | 12 | 20 | 14 | 8.4 | 13 | 11 | 13 | 29 | 8.2 |
| A_0012 | Fumaric acid | 444972 | HMDB0000134 | N.D. | N.D. | 12 | 9.5 | N.D. | N.D. | 11 | 11 | 18 | 9.7 |
| C_0015 | GABA | 119 | HMDB0000112 | 3.9 | 4.8 | 11 | 5.9 | 4.3 | 3.2 | 2.5 | 6.1 | 6.5 | 2.3 |
| A_0087 | GDP | 8977 | HMDB0001201 | 68 | 56 | 70 | 49 | 30 | 57 | 31 | 36 | 97 | 36 |
| C_0056 | Gln | 738 | HMDB0000641 | 354 | 219 | 355 | 312 | 402 | 232 | 365 | 492 | 688 | 340 |
| C_0060 | Glu | 611 | HMDB0000148 | 854 | 737 | 1,193 | 799 | 1,035 | 942 | 942 | 1,176 | 2,102 | 812 |
| A_0052 | Gluconic acid | 10690 | HMDB0000625 | N.D. | N.D. | N.D. | N.D. | N.D. | N.D. | N.D. | N.D. | N.D. | N.D. |
| A_0059 | Glucose 1-phosphate | 65533 | HMDB0001586 | N.D. | 2.4 | 4.7 | 4.2 | 3.3 | 6.7 | 3.8 | 3.7 | 8.0 | 4.3 |
| A_0060 | Glucose 6-phosphate | 5958 | HMDB0001401 | 84 | 60 | 93 | 79 | 70 | 78 | 53 | 65 | 157 | 57 |
| C_0142 | Glutathione (GSH) | 124886 | HMDB0000125 | 83 | 103 | 111 | 87 | 53 | 78 | 36 | 59 | 182 | 68 |
| C_0141 | Glutathione (GSSG)_divalent | 65359 | | 25 | 25 | 36 | 21 | 25 | 43 | 35 | 26 | 69 | 21 |
| C_0003 | Gly | 750 | HMDB0000123 | 190 | 156 | 254 | 167 | 259 | 215 | 176 | 314 | 423 | 167 |
| A_0040 | Glyceraldehyde 3-phosphate | 729 | HMDB0001112 | N.D. | N.D. | N.D. | N.D. | N.D. | N.D. | N.D. | N.D. | N.D. | N.D. |
| A_0041 | Glycerol 3-phosphate | 439162 | HMDB0000126 | 166 | 146 | 254 | 208 | 135 | 236 | 129 | 143 | 403 | 164 |
| A_0003 | Glycolic acid | 757 | HMDB0000115 | N.D. | N.D. | N.D. | N.D. | N.D. | N.D. | N.D. | N.D. | N.D. | N.D. |
| A_0001 | Glyoxylic acid | 760 | HMDB0000119 | N.D. | N.D. | N.D. | N.D. | N.D. | N.D. | N.D. | N.D. | N.D. | N.D. |

### Supplementary Table S1. Quantitative analysis of target metabolites

| ID | Metabolite | PubChem CID | HMDB ID | Concentration (nmol/g) | | | | | | | | | |
|---|---|---|---|---|---|---|---|---|---|---|---|---|---|
| | | | | Control | | | | Lp1534 | | | Lc5805 | | |
| | | | | 0 min | 10 min | 60 min | 300 min | 10 min | 60 min | 300 min | 10 min | 60 min | 300 min |
| | | | | C-1 | C-2 | C-3 | C-4 | Lp-1 | Lp-2 | Lp-3 | Lc-1 | Lc-2 | Lc-3 |
| A_0074 | GMP | 6804 | HMDB0001397 | 23 | 19 | 22 | 17 | 9.7 | 21 | 10 | 14 | 32 | 9.9 |
| A_0097 | GTP | 6830 | HMDB0001273 | 241 | 246 | 324 | 230 | 125 | 217 | 119 | 132 | 410 | 139 |
| C_0065 | Guanine | 764 | HMDB0000132 | 12 | 18 | 18 | 12 | 58 | 21 | 34 | 49 | 37 | 15 |
| C_0136 | Guanosine | 6802 | HMDB0000133 | 66 | 63 | 97 | 72 | 39 | 85 | 65 | 46 | 169 | 64 |
| C_0068 | His | 773 | HMDB0000177 | 66 | 91 | 105 | 60 | 50 | 82 | 37 | 59 | 157 | 45 |
| C_0026 | Homoserine | 12647 | HMDB0000719 | N.D. | N.D. | N.D. | N.D. | N.D. | N.D. | N.D. | N.D. | N.D. | N.D. |
| A_0010 | 2-Hydroxybutyric acid | 440864 | HMDB0000008 | 14 | 12 | 11 | 11 | N.D. | 7.8 | N.D. | N.D. | 12 | 3.0 |
| A_0009 | 3-Hydroxybutyric acid | 441 | HMDB0000011 | 44 | 37 | 44 | 43 | 12 | 29 | 11 | 13 | 60 | 25 |
| C_0039 | Hydroxyproline | 5810 | HMDB0000725 | 6.6 | 5.1 | 5.3 | 2.4 | 4.8 | 5.5 | 3.3 | 8.6 | 29 | 22 |
| C_0049 | Hypoxanthine | 790 | HMDB0000157 | 0.8 | 1.5 | 1.5 | 1.5 | 2.7 | 2.7 | 2.7 | 1.9 | 3.7 | 1.4 |
| C_0043 | Ile | 791 | HMDB0000172 | 185 | 176 | 242 | 151 | 43 | 295 | 39 | 58 | 333 | 93 |
| A_0073 | IMP | 8582 | HMDB0000175 | 4.3 | 5.7 | 6.5 | 4.1 | N.D. | N.D. | 3.1 | 4.3 | 6.6 | 3.5 |
| C_0134 | Inosine | 6021 | HMDB0000195 | 1.7 | 2.7 | 3.4 | 3.3 | N.D. | 3.4 | 2.0 | 2.2 | 5.3 | 2.6 |
| A_0050 | Isocitric acid | 1198 | HMDB0000193 | N.D. | N.D. | N.D. | N.D. | N.D. | N.D. | N.D. | N.D. | N.D. | N.D. |
| A_0007 | Lactic acid | 612 | HMDB0000190 | 464 | 447 | 334 | 269 | 248 | 252 | 134 | 235 | 538 | 157 |
| C_0042 | Leu | 857 | HMDB0000687 | 302 | 234 | 355 | 246 | 62 | 327 | 51 | 87 | 413 | 112 |
| C_0058 | Lys | 866 | HMDB0000182 | 537 | 522 | 673 | 343 | 299 | 582 | 387 | 371 | 1,176 | 407 |
| A_0024 | Malic acid | 525 | HMDB0000156 | 47 | 29 | 46 | 44 | 39 | 45 | 53 | 43 | 79 | 53 |
| A_0084 | Malonyl CoA_divalent | 644066 | | N.D. | N.D. | N.D. | N.D. | N.D. | N.D. | N.D. | N.D. | N.D. | N.D. |
| C_0063 | Met | 876 | HMDB0000696 | 7.3 | 6.4 | 8.9 | 7.6 | 3.5 | 8.2 | 2.8 | 4.9 | 11 | 3.5 |
| A_0103 | NAD$^+$ | 5893 | HMDB0000902 | 27 | 33 | 44 | 29 | 28 | 40 | 28 | 31 | 79 | 24 |
| A_0104 | NADP$^+$ | 5886 | HMDB0000217 | 4.2 | 4.2 | 4.7 | 3.8 | 3.1 | 4.3 | 2.7 | N.D. | 8.0 | 3.3 |
| C_0045 | Ornithine | 389 | HMDB0000214 | 85 | 66 | 121 | 88 | 36 | 70 | 51 | 59 | 197 | 73 |
| A_0029 | 2-Oxoglutaric acid | 51 | HMDB0000208 | N.D. | N.D. | 14 | 8.9 | N.D. | 11 | N.D. | N.D. | 19 | 8.2 |
| A_0013 | 2-Oxoisovaleric acid | 49 | HMDB0000019 | N.D. | N.D. | N.D. | N.D. | N.D. | N.D. | N.D. | N.D. | N.D. | N.D. |
| C_0079 | Phe | 994 | HMDB0000159 | 76 | 67 | 85 | 63 | 52 | 71 | 43 | 71 | 135 | 52 |
| A_0037 | Phosphoenolpyruvic acid | 1005 | HMDB0000263 | 11 | 14 | 14 | 14 | 15 | 22 | 15 | 16 | 24 | 11 |
| A_0048 | 2-Phosphoglyceric acid | 439278 | HMDB0003391 | N.D. | 6.9 | 8.7 | N.D. | N.D. | 8.5 | 5.8 | N.D. | 10 | 5.4 |
| A_0047 | 3-Phosphoglyceric acid | 439183 | HMDB0000807 | 48 | 56 | 73 | 48 | 52 | 73 | 72 | 70 | 100 | 44 |
| A_0063 | 6-Phosphogluconic acid | 91493 | HMDB0001316 | 4.9 | 4.3 | 4.4 | 3.4 | N.D. | 3.1 | N.D. | N.D. | 9.3 | 3.3 |
| C_0023 | Pro | 614 | HMDB0000162 | 121 | 159 | 164 | 107 | 98 | 104 | 74 | 118 | 283 | 119 |
| A_0076 | PRPP | 7339 | HMDB0000280 | N.D. | N.D. | N.D. | N.D. | N.D. | N.D. | N.D. | N.D. | N.D. | N.D. |
| C_0006 | Putrescine | 1045 | HMDB0001414 | 167 | 210 | 255 | 184 | 182 | 208 | 144 | 208 | 337 | 145 |
| A_0004 | Pyruvic acid | 1060 | HMDB0000243 | N.D. | N.D. | N.D. | N.D. | N.D. | N.D. | N.D. | N.D. | N.D. | N.D. |
| A_0058 | Ribose 5-phosphate | 439167 | HMDB0001548 | N.D. | N.D. | N.D. | N.D. | N.D. | N.D. | N.D. | N.D. | N.D. | N.D. |
| A_0057 | Ribulose 5-phosphate | 439184 | HMDB0000618 | N.D. | 2.7 | N.D. | 4.1 | N.D. | 4.2 | 3.3 | N.D. | 8.7 | 2.7 |
| C_0148 | *S*-Adenosylmethionine | 34755 | HMDB0001185 | 12 | 12 | 17 | 14 | 3.2 | 13 | 4.4 | 3.6 | 24 | 6.9 |
| C_0008 | Sarcosine | 1088 | HMDB0000271 | N.D. | N.D. | N.D. | N.D. | N.D. | N.D. | N.D. | N.D. | N.D. | N.D. |
| A_0064 | Sedoheptulose 7-phosphate | 165007 | HMDB0001068 | 15 | 11 | 23 | 15 | 9.7 | 15 | 9.8 | 7.9 | 27 | 11 |
| C_0017 | Ser | 617 | HMDB0000187 | 276 | 262 | 412 | 242 | 316 | 379 | 308 | 349 | 719 | 251 |
| C_0055 | Spermidine | 1102 | HMDB0001257 | 55 | 58 | 74 | 42 | 57 | 67 | 52 | 71 | 143 | 62 |
| C_0099 | Spermine | 1103 | HMDB0001256 | N.D. | N.D. | N.D. | N.D. | N.D. | N.D. | N.D. | N.D. | N.D. | N.D. |
| A_0014 | Succinic acid | 1110 | HMDB0000254 | 124 | 81 | 130 | 119 | 60 | 115 | 47 | 89 | 159 | 100 |
| C_0027 | Thr | 6288 | HMDB0000167 | 238 | 223 | 321 | 239 | 152 | 254 | 121 | 190 | 501 | 192 |
| C_0116 | Thymidine | 5789 | HMDB0000273 | N.D. | N.D. | N.D. | N.D. | N.D. | N.D. | N.D. | N.D. | N.D. | N.D. |
| C_0033 | Thymine | 1135 | HMDB0000262 | N.D. | N.D. | N.D. | N.D. | N.D. | N.D. | N.D. | N.D. | N.D. | N.D. |
| C_0101 | Trp | 1148 | HMDB0000929 | 31 | 26 | 36 | 27 | 16 | 34 | 14 | 19 | 54 | 17 |
| C_0085 | Tyr | 1153 | HMDB0000158 | 57 | 55 | 75 | 50 | 40 | 53 | 29 | 50 | 91 | 28 |
| C_0051 | Tyramine | 5610 | HMDB0000306 | 1.6 | 1.5 | 1.6 | 1.9 | N.D. | 2.0 | 0.4 | N.D. | 2.1 | 1.7 |
| A_0079 | UDP | 6031 | HMDB0000295 | 14 | 14 | 18 | 13 | 12 | 17 | 11 | 13 | 32 | 13 |
| A_0068 | UMP | 6030 | HMDB0000288 | 11 | 8.5 | 11 | 11 | 4.3 | 11 | 8.1 | 9.8 | 18 | 8.4 |
| C_0021 | Uracil | 1174 | HMDB0000300 | N.D. | N.D. | N.D. | N.D. | N.D. | 7.8 | N.D. | N.D. | 12 | N.D. |
| C_0118 | Uridine | 6029 | HMDB0000296 | 5.1 | 5.0 | 6.0 | 8.4 | N.D. | 7.3 | 3.8 | 5.3 | 12 | 7.2 |
| A_0092 | UTP | 6133 | HMDB0000285 | 30 | 32 | 50 | 34 | 27 | 35 | 23 | 26 | 72 | 24 |
| C_0024 | Val | 1182 | HMDB0000883 | 303 | 299 | 380 | 255 | 85 | 441 | 72 | 117 | 484 | 146 |
| C_0009 | β-Ala | 239 | HMDB0000056 | 67 | 58 | 91 | 68 | 44 | 47 | 22 | 48 | 105 | 40 |

IDs consist of a prefix indicating the measurement mode followed by a sequential number, where "C" and "A" denote cation mode and anion mode, respectively.

N.D., Not Detected (target for analysis, but below the limit of detection)

N.A., Not Available (target for calculation, but could not be calculated due to insufficient data)

Compounds are sorted in ascending alphabetical order by compound name.
